# Wearing an Eye Mask During Overnight Sleep Improves Episodic Learning and Alertness

**DOI:** 10.1101/2022.01.20.477083

**Authors:** Viviana Greco, Damiana Bergamo, Paola Cuoccio, Karen R. Konkoly, Kike Muñoz Lombardo, Penelope A. Lewis

**Affiliations:** Cardiff University Brain Research Imaging Centre (CUBRIC), School of Psychology, Cardiff University, Maindy Rd, Cardiff, CF24 4HQ, UK; IMT School for Advanced Studies Lucca, 55100 Lucca, Italy; Department of Psychology, University of Padua, Padova, 35122, Italy; Department of Psychology, Northwestern University, Chicago, USA

**Keywords:** Sleep, eye mask, learning, episodic memory, alertness

## Abstract

Ambient light can influence sleep structure and timing. We explored how wearing an eye-mask to block light during overnight sleep impacts on memory and alertness, changes that could benefit everyday tasks like studying or driving. In Experiment 1, ninety-four 18–35-year-olds wore an eye-mask while they slept every night for a week and underwent a control condition in which light was not blocked for another week. Five habituation nights were followed by a cognitive battery on the sixth and seventh days. This revealed superior episodic encoding and an improvement on alertness when using the mask. In Experiment 2, thirty-five 18–35-year-olds used a wearable device to monitor sleep with and without the mask. This replicated the encoding benefit and showed that it was predicted by time spent in slow wave sleep. Our findings suggest that wearing an eye-mask during overnight sleep can improve episodic encoding and alertness the next day.

**Statement of relevance:** Sleep is crucial for alertness and for preparing the human brain to encode new information. However, it can be disrupted by external stimuli such as light or sounds. This study explored wearing an eye mask as a potential cognitive enhancer which protects overnight sleep by blocking ambient light. We found that wearing a mask increased alertness and facilitated the encoding of novel information the next day. Furthermore, the benefit to memory was predicted by time spent in slow wave sleep while wearing the mask. This suggests wearing an eye mask during sleep is an effective, economical, and non-invasive behaviour that could benefit cognitive function and lead to measurable impacts on every-day life.

## Introduction

Sleep plays an essential role in many physiological functions including immune control, energy conservation, homeostatic restoration and memory processing (Diekelmann & Born, 2010; Killgore, 2010; Lange et al., 2010). Sleep quality and quantity is crucial for brain function, and studies have indicated that a night of sleep deprivation or multiple nights of partial sleep restriction have a negative impact on cognition, for instance affecting subsequent memory encoding (Cousins et al., 2018; van der Werf et al., 2009, 2011; Yoo et al., 2007). Moreover, optimal academic performance is strongly related to the timing, quality, and quantity of sleep (Curcio et al., 2006).

In mammals, the sleep-wake cycle is regulated by the suprachiasmatic nuclei (SCN) of the anterior hypothalamus (Daan et al., 1984). SCN activity is strongly synchronised by the light-dark cycle via intrinsically photosensitive retinal ganglion cells (Blume et al., 2019; Wams et al., 2017). The tight interaction between light and sleep regulation is therefore clear, with a large body of evidence supporting the impact of light on sleep timing, macro-architecture and duration (Badia et al., 1991; Borbely, 1982; Borbély et al., 2016; Dijk & Archer, 2009; Schmidt et al., 2011; Wams et al., 2017). A recent study conducted by Wams and colleagues assessing the link between light exposure and subsequent sleep, revealed that subjects with earlier exposure to light spent significantly more time in slow wave sleep (SWS) at the expense of rapid-eye movement (REM) sleep (Wams et al,2017).

Non-pharmacological methods for improving sleep are a topic of great current interest (Wunderlin et al., 2021), and the use of an eye mask to prevent light from reaching the retina during overnight sleep has been demonstrated to positively affect subjective sleep quality in intensive care patients who are systematically exposed to high levels of light (Bani Younis et al., 2019; Locihová et al., 2018). In the current study, we set out to investigate the benefits of wearing an eye mask to block light during normal sleep in the home. Declarative learning (Van Der Werf et al., 2009) and vigilant attention (Anderson et al., 2010; Van Der Werf et al., 2011) are both known to be sleep sensitive. Given the practical importance of these abilities in everyday life, for instance in studying at school or in driving a car (Dorrian et al., 2019), we wanted to examine the impact of the eye mask manipulation of these abilities. With this aim in mind, we ran a within-subject design to look at these cognitive processes (Experiment 1) and a follow-up study which examined sleep architecture (Experiment 2).

## Method

### Participants

All participants were healthy volunteers, with no history of drug/alcohol abuse, psychological, neurological, or sleep disorders. We selected participants who reported no hypersensitive skin or contact allergies and no problems falling asleep with open shutters and wearing both an eye mask and a wearable EEG device (Dreem headband, DH, (Arnal et al., 2020)). Participants agreed to abstain from alcohol and caffeine throughout the experiment. Additionally, the online screening ensured that they had not worn an eye mask for sleep before and they agreed not to nap on the days of the experiment.

The sample size of Experiment 1 was determined using a power calculation based on a pilot study (n=8) on the Paired associate learning (PAL) task and based on a paired sample t-test (G*Power Version 3.1.9.6; Erdfelder et al., 2009). This pilot predicted 80% power to detect a medium-size effect (Cohen’s d=0.3) with 88 subjects and a conventional α of 0.05. 94 native English speakers (59F, age range: 18-35, M=21.07, SD=2.74) took part in the study. Of these, five were excluded due to voluntary withdrawal, so our final dataset included 89 participants (54F, M=20.98, SD=2.68). Due to technical failures when executing the tasks, a further six participants were excluded from the PAL, four were excluded from the Psychomotor Vigilance Test (PVT) and three from the Motor skill learning (MSL) analysis.

Experiment 2 was undertaken by thirty-seven native English and Italian speakers (29F, age range: 18-35, M=23.03, SD=3.52). Of these, 4 were excluded due to difficulty falling asleep before 5am (n=1), sudden notification of working commitments (n=1) and voluntary withdrawal (n=2). Our final dataset therefore included 33 participants (25F, M=23.09, SD=3.57). Sample size was predetermined using a formal power analysis for correlation analysis. A sample size of 29 was needed to detect a correlation coefficient of 0.5 with a conventional α of 0.05 and 80% power. The number recruited slightly exceeded the number needed. All participants gave informed consent for the experiments, which were approved by the Ethics Committee of the School of Psychology at Cardiff University and received monetary compensation for their participation.

### Procedure

The study design of both experiments is outlined in Figure 1.

**Fig. 1.**
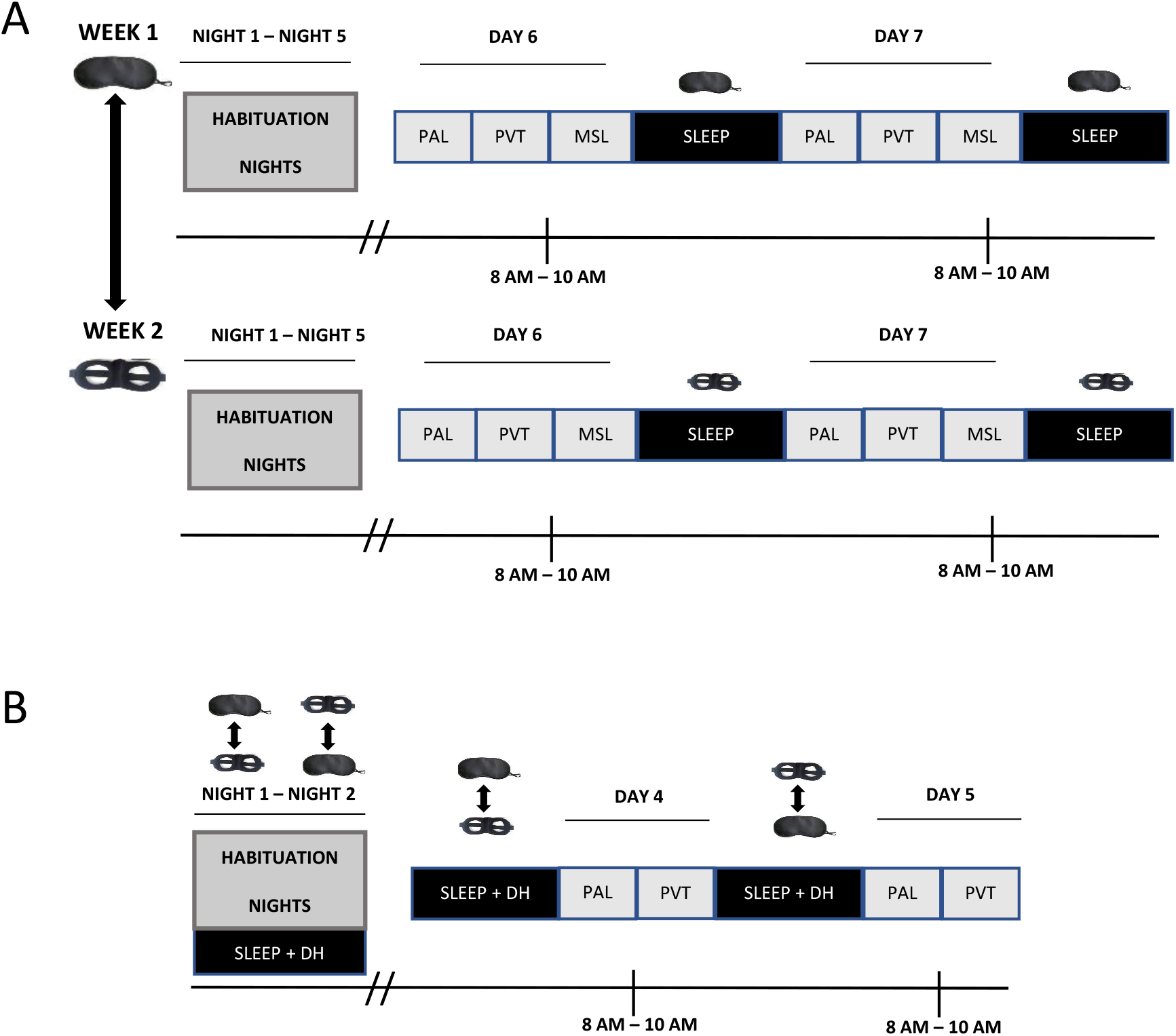
Experimental procedure. **A.** Experiment 1 consisted of 2 consecutive weeks in which, in a counterbalanced order, ambient light was blocked with an eye mask during sleep for one week, or not blocked with a control mask for the other week. Night1-Night5: participants slept at home wearing a mask (eye mask or control). Day6-Day7: participants performed the Paired associate learning (PAL), the Psychomotor vigilance test (PVT) and the Motor-skill learning (MSL) task. **B.** Experiment 2 consisted of 5 days, 2 habituation nights and 2 experimental days. For the entire study duration, participants slept with an eye mask or a control mask (counterbalanced order) together with the Dreem headband (DH). In the morning of Day4-Day5, participants completed the PAL and the PVT.

Experiment 1 was conducted over two summers (end of June – end of September) of 2018 and 2019. We chose the summer months because we suspected that the eye mask would be more helpful when dawn occurred early (as early as 5am at midsummer in Cardiff, UK). The 2018 study involved two consecutive weeks in counterbalanced order, in which ambient light was blocked with an eye mask during sleep for the experimental week, and not blocked for the control week. Each week consisted of five habituation nights, followed by two consecutive testing days (Day6 and Day7 respectively). During the habituation nights, participants were instructed to sleep at home (wearing eye mask or control) and to maintain their regular sleep-wake habits. On Day6 and 7, participants arrived in the sleep laboratory at Cardiff University Brain Research Centre (CUBRIC) at 8am-10am and performed three cognitive tasks: PAL, PVT, MSL. The 2019 study was identical to the 2018 study, except that in the control condition participants wore a modified eye mask with two big holes cut over the eyes, such that it did not block the light. This was done in order to control for any discomfort caused by wearing a mask. Computer-based tasks were executed using Matlab 2016 or 2017 (The MathWorks Inc., Natick, MA, USA).

Experiment 2 was conducted over the summer of 2020 and consisted of four nights (two habituation and two experimental), in which participants were asked to sleep at home with the DH and an eye mask or modified mask with holes, in a counterbalanced order. The first two habituation nights were used to accustom participants to sleeping while wearing both the DH and one of the masks. Between 8 and 10 am of Days 4 and 5, subjects performed the learning part of the PAL and the PVT, see Supplemental Method for details of the tasks. Tasks were executed online using PsychoPy3 Experiment Runner (v2020.1.3, (Peirce et al., 2019)). From now on, we will refer to the two conditions as: “eye mask”, for the normal sleep mask, and “control”, for both no mask (2018 participants) and the modified mask with holes over the eyes (2019/2020 participants).

In both experiments, an online sleep diary was completed every morning and subjective alertness, was measured with the Stanford Sleepiness Scale (SSS, (Hoddes et al., 1973)) on the morning of each experimental day. Participants were asked to sleep with open shutters/curtains for the entire duration of the study.

### Wearable EEG device: Dreem headband

In Experiment 2 sleep macro-architecture was recorded using the DH that automatically records, stores and analyses physiological data (Arnal et al., 2020). The DH consists of five dry-EEG electrodes (O1, O2, FpZ, F7, F8). The signal is recorded with a sampling frequency of 250 Hz with a 0.4-35 Hz bandpass filter. The DH allowed participants to sleep in their own environment rather than in a laboratory setting, increasing sleep quality and comfort levels. Recent validation of the automatic sleep stage classification of the DH, compared to the standard polysomnography (PSG), showed that the automatic algorithm can reliably perform sleep staging (Arnal et al., 2020).

We examined the time spent in each sleep stage during the two experimental nights (nights 3 and 4). Relationships between behavioural measures and sleep were assessed with Pearson’s correlations, or Spearman’s Rho if Shapiro-Wilk tests indicated a non-normal distribution. All statistical tests were 2-tailed and considered significant at p<0.05. Analyses were conducted in R (version 4.0.2, R Core Team, 2020). Results are presented as mean ±SEM. Four participants didn’t start the DH recording correctly, so sleep data was not collected. Sleep macrostructure analysis were therefore based on N=29 participants.

### Behavioural analysis

We implemented a linear mixed-effects (LME) analysis in R with the “lme4” package (Bates et al., 2012). In all models, we included a fixed effect factor related to the type of the mask (two levels: eye mask, control mask) and participant IDs as random effects. In Experiment 1 we also included a random intercept for the year of the experiment (2018, 2019): lmer (DV~‘Mask_type’+ (1|Subject) + (1|Year), data, REML=FALSE), where DV is the dependent measure. Note that we included ‘year’ in order to capture any variability due to slight differences in the control condition in 2018 and 2019. Predictor was coded as follows: ‘Mask_type’: eye mask=1, control=0. To determine statistical significance, we conducted a likelihood ratio test (LRT) in which the full model with fixed and random effects was compared to a reduced model with random effects only. Chi-square statistic (χ^2^), associated degrees of freedom and p-values for the final models are provided. Additionally, Akaike Information Criteria (AIC) related to a full model and to a reduced model, was also reported. Significance threshold was set at 0.05. For all models, visual inspection of residual plots was used to assess the model assumptions of linearity, homoscedasticity and normality of the residuals. Effect sizes were computed using “lsr” package (Navarro, 2015). All figures were created using ggplot2 R-package (Wickham, 2009). Descriptive statistic of the tasks for both experiments is reported in Supplemental Results (Experiment 2), Table S1.

### Questionnaires

The SSS was used to provide a subjective indication of sleepiness, with participants rating their current state on a 7-point Likert scale, where 1 is most alert and 7 is least alert (Hoddes et al., 1973).

A sleep diary was used to gather information about units of alcohol and caffeine consumed, sleep duration and the regularity of sleep-wake cycle. In Experiment 2 we also assessed the comfort of the masks and the DH on a five-point Likert scale (1: “Very uncomfortable” to 5: “Very comfortable”). Likewise, self-rating of sleep quality was measured on a Likert scale (1: “Very poor” to 5: “Very good”). Paired-sample t-tests were used to evaluate whether sleep quality differed after the use of the eye mask or the control. When the assumption of normality was violated, non-parametric Wilcoxon signed-rank tests were conducted instead. All tests were two-tailed and a *p*-value < 0.05 was used for all analyses. Effect sizes were computed using “rcompanion” and “lsr” packages (Mangiafico, 2020; Navarro, 2015).

## Results

### Experiment 1

#### Paired associate learning task (PAL)

We first assess whether wearing the eye mask affected learning performance on the word-pair associate task. Inclusion of ‘Mask_type’ in our LME significantly improved model fit (χ^2^_1_ = 5.21; *p* = 0.022), showing significantly better learning after wearing the eye mask compared to the control (eye mask: 65.06±0.69 vs control: 63.87±0.67; b= −1.19, *p* = 0.023, d= 0.19; Figure 2a, Table S2). An additional LME model which we fit to examine whether our experimental intervention had an impact on overnight declarative memory consolidation, revealed no effect of the eye mask on overnight change in memory performance (Table S2).

**Fig. 2.**
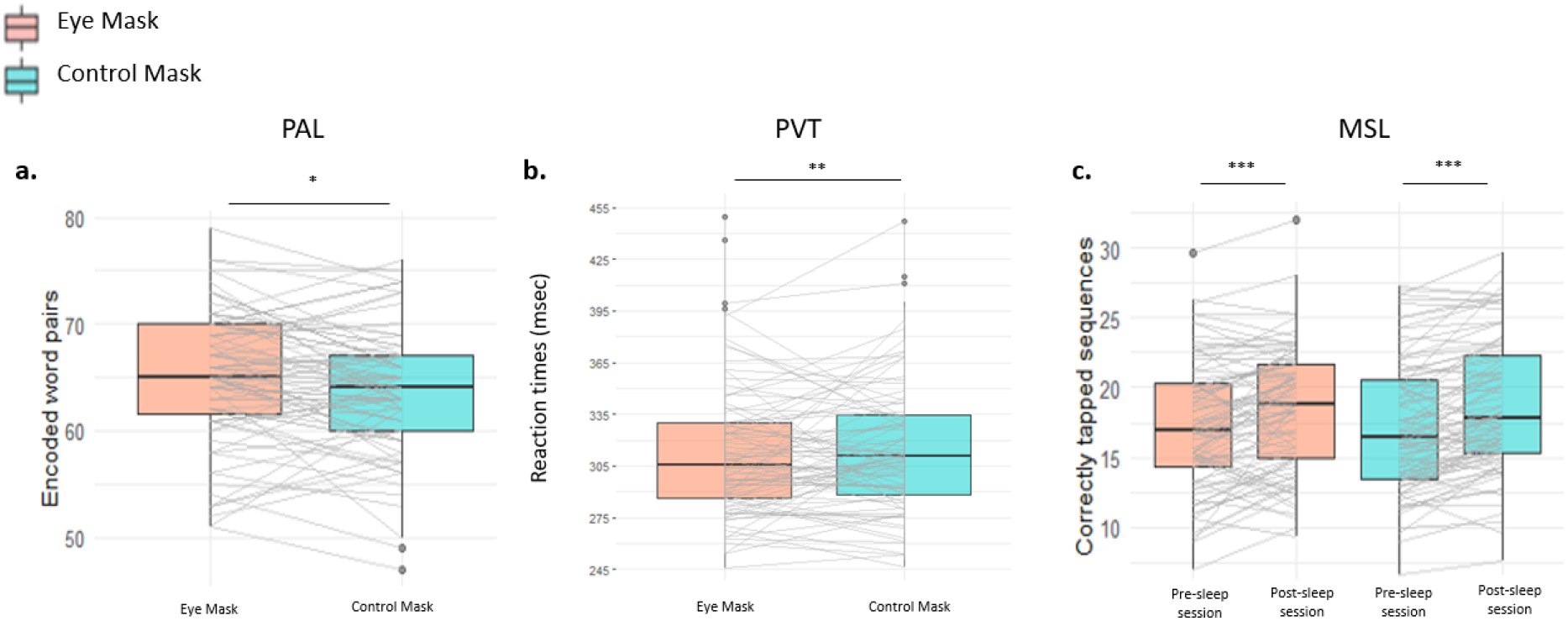
Behavioural results. Boxplots for **(a)** learning performance on the Paired associate learning task (PAL), **(b)** reaction times on the Psychomotor Vigilance Test (PVT) **(c)** number of correctly tapped sequences on the Motor Skill Learning task (MSL). Boxplots show the median (inner thick horizontal line), the 25^th^ and 75^th^ percentiles (the bottom and the top of the box), the largest value within 1.5 * interquartile range (upper whisker) and the smallest value within 1.5 * interquartile range (lower whisker). Data beyond the end of the whiskers are outliers. *p< 0.05, **p<0.01, ***p<0.001. ns: non-significant.

#### Psychomotor vigilance test (PVT)

We next assessed the impact of wearing the eye mask on the psychomotor vigilance task. PVT responses from the two testing days were combined in an LME which tested for differences in reaction time after wearing the eye mask as compared to the control. This revealed that blocking ambient light impacted significantly upon reaction times (χ^2^_1_ = 7.32; *p* = 0.006), with participants responding faster after wearing the eye mask (eye mask: 310.26±2.79 vs control: 316.37±2.94; b= 316.37, *p* = 0.006, d= 0.16; Figure 2b, Table S2).

#### Motor-skill learning (MSL)

We examined motor-skill learning by fitting a linear mixed effects model to the number of correctly tapped sequences with fixed effects of ‘Mask_type’ (eye mask and control), Day (Day6 and Day7), and their interaction with random effects for participants and for the year of the experiment (2018 and 2019).

This showed no effect of mask (b= −0.00, *p* = 1.000), but a main effect of Day (b= 1.19, *p* = 0.000), indicating that our participants performed the task faster after sleep irrespective of mask or control condition. There was no interaction between ‘Mask_type’ and Day (b= 0.24, *p* = 0.597), indicating that our intervention did not modulate performance on this task (Figure 2c). Examination of the absolute overnight change in performance revealed no differences between the two type of masks (Table S2).

#### Questionnaires

The LRT on the SSS performed on both testing days combined, revealed no effect of mask on subjective alertness (χ^2^_1_ = 3.11, *p* = 0.078; eye mask: 2.22±0.06 vs control: 2.34±0.06). Moreover, the eye mask had no impact on the actual number of hours which participants reported sleeping since a Wilcoxon-signed rank test on the total number of hours slept across the week (as indexed by the sleep diary) revealed no differences between conditions (eye mask: 8.24±0.09 vs control: 8.26±0.11, Z= −0.27, *p*= 0.785; N= 79). Notably, ten participants were excluded from this analysis due to the poor compliance in completing the sleep diary.

### Experiment 2

In Experiment 2 we sought to build on Experiment 1 by adding objective measurements of time spent in each sleep stage through the use of the Dreem headband. However, the eye mask manipulation did not cause any changes in the sleep macrostructure, as reported in Table S3. For consistency, we collected similar behavioural and questionnaire data as in Experiment 1. This leads to a significant result in the PAL, reported below, but no other significant findings (see Supplemental Results (Experiment 2)).

#### Paired associate learning task (PAL)

In our assessment of mask vs control on word-pair encoding, the LRT revealed that the inclusion of ‘Mask_type’ in the model provided a better fit for the data (χ^2^_1_ = 3.91; *p* = 0.047). Thus, in keeping with Experiment 1, learning performance was better after wearing the eye mask than the control (eye mask: 69.9±1.89 vs control: 67.7±1.80; b= 2.18, *p*= 0.049, d= 0.22; Figure 3a, Table S4). Notably, this result did not change when 4 outliers were removed (eye mask: 72.1±1.22 vs control: 69.8±1.22; b= 2.35, *p* = 0.030, d= 0.38; Table S4, see Supplemental Results (Experiment 2) for details).

**Fig 3.**
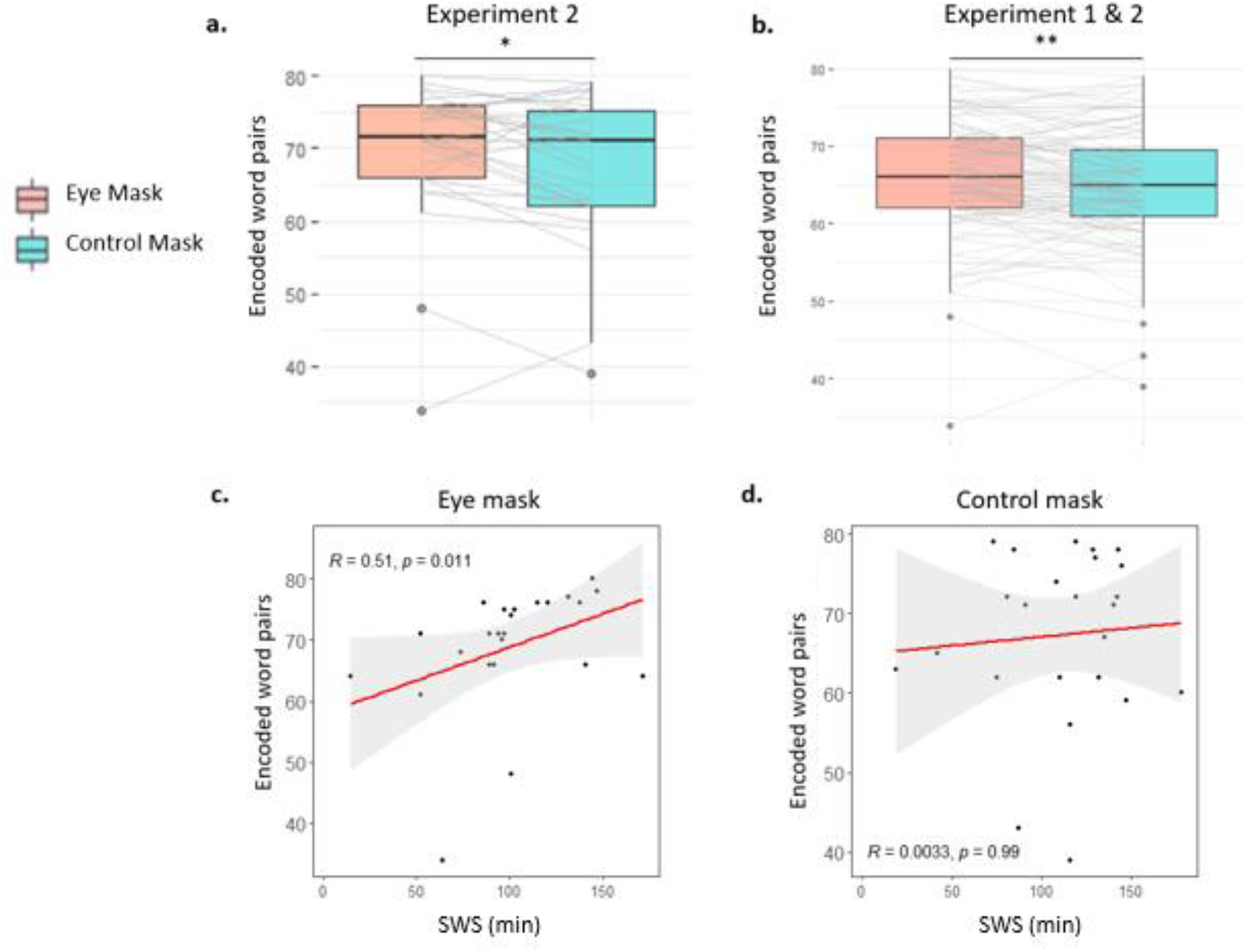
**a:** Experiment 2. Paired associate learning (PAL) results (N=28). Boxplots for learning performance on the PAL after a night of sleep wearing the eye mask (pink) or the control mask (light blue). **b:** Combined results of the encoding performance on the PAL from Experiment 1 and 2 (N=112). **p<0.01; *p< 0.05. **c:** Significant Spearman’s (rank) correlation between the time spent in SWS (minutes) and the learning perfomance on the word-pairs after a night wearing the eye mask. Note that when N=3 outliers were removed, the correlation was still significant (r_s_=0.44, p=0.04). **d:** Spearman’s (rank) correlation between time spent in SWS (minutes) and learning perfomance on the word-pairs after a night wearing the control mask.

To examine the overall dataset, we combined the PAL results related to the learning performance from both Experiment 1 and Experiment 2 (N=112): lmer (DV~ ‘Mask_type’ + (1|Subject) + (1|Year:2018-2019-2020), data, REML=FALSE). This indicated that the inclusion of the eye mask effect improved model fit (χ^2^_1_ = 9.01; *p* = 0.002). Overall, the use of the eye mask had a positive effect upon subsequent learning (eye mask: 66.6±1.5 vs control: 65.1±1.5; b=1.44, *p* = 0.002, d= 0.19; Table S4, Figure 3b). Given these results, we conclude that wearing the eye mask was beneficial for declarative memory encoding the next day.

In light of previous studies demonstrating that SWS plays a major role for subsequent encoding (Antonenko et al., 2013; van der Werf et al., 2009, 2011; Yoo et al., 2007), we tested for correlations between the learning of hippocampus-dependent memory and SWS time. Learning performance after a night wearing the eye mask was positively correlated with SWS time (r_s_= 0.51, *p* = 0.011; Figure 3c), whereas there was no such correlation after a night wearing the control mask (r_s_= 0.00, *p* = 0.99; Figure 3d). No other correlations were significant (all *p* > 0.05).

#### Questionnaires

Sleep diary data revealed no differences in the number of hours slept while wearing the eye mask (7.15±16.66) or the control mask (7.18±16.82; t(23)= −0.11, *p*= 0.914, N= 24). Likewise, there was no significant difference in self-rating of sleep quality (eye mask: 3.13±0.19 vs control: 2.84±0.16; Z= −1.53, *p* = 0.131, N= 31). Participants rated the control mask as more uncomfortable than the eye mask (eye mask: 3.10±0.17 vs control: 2.40±0.12; Z= −2.86, *p* = 0.005, N= 30), but this did not impact on the comfort of the DH with mask (eye mask: 2.83±0.18 vs control: 2.83±0.17; Z=0.00, *p* = 1.000, N= 30).

## Discussion

Our results demonstrate that wearing an eye mask during overnight sleep can facilitate both new learning and alertness the next day.

Sleep before learning has been shown to impact upon subsequent encoding (Cousins et al., 2018; McDermott et al., 2003; van der Werf et al., 2009, 2011; Yoo et al., 2007). For instance, Van Der Werf and colleagues revealed impaired declarative encoding accompanied by a decreased hippocampal activation after selective deprivation of SWS (van der Werf et al., 2009). Contrary to previous studies demonstrating a reduction in the total amount of REM sleep in response to bright morning light exposure (Dijk et al., 1987, 1989), our examination of sleep macrostructure did not reveal any differences between a night spent wearing an eye mask and a night spent wearing a control mask. Interestingly, however, the memory improvement associated with the eye mask was positively correlated with the time spent in SWS. Encoding of declarative materials has been shown to be enhanced when slow wave activity (SWA) is artificially increased through transcranial slow oscillation stimulation (Antonenko et al., 2013). The synaptic homeostasis hypothesis (SHY) posits that SWA (0.5-4 Hz), a hallmark of SWS, promotes the global down-scaling of synapses that have become saturated during preceding periods of wakefulness and thus restores capacity for the encoding of new information (Tononi & Cirelli, 2003, 2014). Given this literature, we speculate that while wearing an eye mask did not increase time spent in SWS it may have increased SWA. However, we were unable to measure SWA due to the minimal nature of recording from the DH.

Turning to vigilance, the current findings suggest that wearing an eye mask has a beneficial effect upon behavioural alertness. To be specific, after a night of sleep spent wearing the eye mask, participants responded faster in the PVT. The PVT is among the most widely used measure of behavioural alertness and sustained attention, with negligible practice and aptitude effects over repeated administrations (Basner & Dinges, 2011; Lim & Dinges, 2008). A variety of studies investigating the effect of light exposure on sustained attention have used this measure after exposing participants to light treatment after a period of sleep or sleep deprivation, with the aim of counteracting sleep deprivation (Phipps-Nelson et al., 2003; Münch et al., 2016; Comtet et al., 2019). Our observation that PVT is improved after wearing an eye mask merits consideration because of the crucial role played by behavioural alertness in many real-world tasks, ranging from driving to other activities that require rapid responses (Dorrian et al., 2019), and because of the ecological setting in which our study was conducted. In fact, our participants slept in the comfort of their own home and were not sleep deprived; moreover, no manipulation of natural light was applied.

Turning to the motor skill learning task, consistent with the literature, we demonstrated that subjects significantly improved on this task after a retention period of sleep (Walker et al., 2002). This overnight learning gain has previously been shown to correlate with the amount of Stage2 sleep obtained, particularly in the last quarter of the night (Walker et al., 2002). However, despite this overnight enhancement, the use of an eye mask does not appear to provide further benefit to this task.

Subjective sleep quality, as assessed with the sleep diary, showed no benefits of the eye mask in either experiment. It deserves mention that even though participants in Experiment 2 reported that sleeping with the control mask was more uncomfortable in comparison with the eye mask, this did not impact self-reported sleep quality, morning alertness, or sleep parameters.

Overall, our findings suggest that a simple manipulation – the use of an eye mask during sleep - can lead to superior memory performance and higher alertness the next day. These findings have broad implications for performance of the many daytime tasks that require learning in educational and cultural contexts, in which particularly effective encoding will determine opportunities for growth, as well as a fast response to external stimuli. Given the current climate of life-hacking, sleep monitoring, and cognitive enhancers, our findings suggest the eye mask as a simple, economical, and non-invasive way to get more out of a night of sleep.

## Notes

### Competing Interest Statement

The authors have declared no competing interest.

