## Supplemental material for "Wearing an Eye Mask During Overnight Sleep Improves Episodic Learning and Alertness"

### **Supplemental Online Material**

#### **Supplemental Method**

***Paired associate learning task (PAL)***

Declarative memory was assessed with a paired associate learning task (PAL). On the first testing day (Day6 of both weeks), participants were first instructed to learn 80 semantically related pairs of English nouns presented on a screen for 3500ms with an interstimulus interval (ISI) of 1000ms. We had two different lists of word-pairs, counterbalanced between subjects. The learning session was followed by an immediate cued recall test, in which subjects were presented with the first noun of every pair in a random order and asked to recall the associated noun by typing it into a computer keyboard. Unlimited time was given to type an answer, and word-pairs were repeatedly presented until the subject reached 60% accuracy. After each response, the correct answer was displayed for 2s, allowing the subject to relearn the pair if necessary. Followed by 10minutes of Psychomotor Vigilance Test (PVT), a final cued recall without feedback was presented and used to assess participants’ pre-sleep score. Approximately 24h later, a delayed recall without feedback took place and it was used to calculate the post-sleep score. We calculated: (i) learning performance as the number of correctly recalled pairs in the final cued recall (pre-sleep score); (ii) absolute consolidation performance as the number of correctly recalled words in the delayed recall minus the final cued recall (post-sleep score – pre-sleep score).

***Psychomotor Vigilance Test (PVT)***

A 10minutes PVT performed on both testing days, required to focus on a fixation cross and to respond (and stop by pressing the space bar on the keyboard), as quickly as possible, to a millisecond counter that randomly appeared between 2-10s. When the counter was stopped, reaction times (in ms) remained on the screen for 1s and served as a feedback. If the space key was not pressed within 2000ms, an onscreen message reminded participants to pay attention. Responses below 100ms (false starts) and above 500ms (attention lapses) were excluded from the analysis before the average score from the remaining trials was calculated (Basner & Dinges, 2011).

***Motor-skill learning task (MSL)***

The Motor-skill learning task (MSL) (Walker et al., 2002) required subjects to repeatedly tap, with their non-dominant hand, a fixed five-digit sequence (e.g. 4-1-3-2-4) pressing four numeric keys on a computer keyboard, as quickly and accurately as possible. On the first testing day of both weeks (Day6), the pre-sleep session required to repeat the numeric sequence throughout 12 blocks of trials of 30s each, with a 30s ISI. A short practise round of 30s was performed before beginning the task. On the second testing day of both weeks (Day7), the post-sleep session followed the same procedure, but 3 blocks of 30s had a different control sequence at the end, to allow determination of whether the overnight improvement was sequence specific. We used two sequences for the present task, which order was balanced across subjects. Pre-sleep performance was calculated by averaging correctly tapped sequences in the last three blocks, while post-sleep performance was assessed by averaging the first three blocks. Performance improvement was calculated as the difference between pre-sleep and post-sleep scores (absolute overnight change).

**Supplemental Results (Experiment 2)**

Experiment 2 was conducted in order to look at the impact of the eye mask manipulation on sleep parameters. For consistency, we also collected PAL, PVT and questionnaire data as in Experiment 1. However, because a power calc for PVT based on the results of Experiment 1 suggested an n of approximately 390 for 80% power, and this was not a feasible target for an experiment using the Dreem headbands, we did not expect a significant result for this task.

***Paired associate learning (PAL)***

In our PAL analysis (see main text), four datapoints that were more than 1.5 inter-quartile range (IQR) below the first quartile or above the third quartile were detected as outliers. To test whether these outliers influenced our result we removed them and re-ran the analysis. When the LME model was performed without these outliers, the LRT revealed that the inclusion of the ‘Mask_type’ in the model provided a better fit for the data compared to a model without it (χ^2^_1_=4.750; *p*=0.029, TableS4). A significantly greater performance at learning was found after wearing the eye mask than the control (eye-mask: 72.1±1.22 vs control: 69.8±1.22; p=0.033, d=0.633) (FigureS1). Thus, removal of the outliers did not change our result.

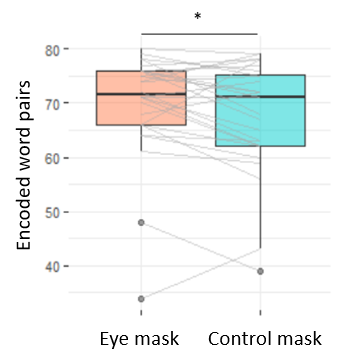

**Fig. S1**. Paired associate learning (PAL) results from Experiment 2 without N=4 outliers. *p < 0.05.

***Psychomotor Vigilance Test (PVT)***

When we examined sustained attention, the LRT revealed no significant effect of the mask (χ^2^_1_=1.82; *p*=0.176), suggesting no specific effect of the eye mask on reaction times (b=-7.33, *p*=0.18; TableS4). Furthermore, analysis of the association between sleep parameters and measure of behavioural alertness revealed no significant correlation (all *p*>0.05). Notably, the experiment was severely underpowered for examination of the PVT.

***Questionnaires***

With regards to the ratings of sleepiness, the LRT showed no significant difference between the models (χ^2^_1_=0.90; *p*=0.342), suggesting that the eye mask had no effect on the SSS relative to the control (eye-mask: 3.57±0.24 vs control: 3.36±0.25; b=0.21, *p*=0.346).

Table S1.

*Descriptive statistic of the experimental tasks.*

| **EXPERIMENT 1** | | | | | |
| --- | --- | --- | --- | --- | --- |
| **DAY 6** | | | | | |
|  |  | **Mean** | **Median** | **95% CI** | **SEM** |
| **PAL** | **Eye mask** | 65.06 | 65.00 | 63.69 – 66.43 | 0.69 |
|  | **Control mask** | 63.87 | 64.00 | 62.53 – 65.20 | 0.67 |
| **PVT** | **Eye mask** | 311.58 | 307.87 | 303.28 – 319.87 | 4.17 |
|  | **Control mask** | 314.13 | 310.00 | 306.06 – 322.20 | 4.06 |
| **MSL** | **Eye mask** | 17.27 | 17.00 | 16.27 – 18.26 | 0.50 |
|  | **Control mask** | 17.27 | 16.50 | 16.25 – 18.29 | 0.51 |
| **DAY 7** |  |  |  |  |  |
| **PAL** | **Eye mask** | 63.73 | 63.00 | 62.32 – 65.15 | 0.71 |
|  | **Control mask** | 62.97 | 64.00 | 61.48 – 64.47 | 0.75 |
| **PVT** | **Eye mask** | 308.95 | 303.20 | 301.51 – 316.39 | 3.74 |
|  | **Control mask** | 318.60 | 313.46 | 310.11 – 327.10 | 4.27 |
| **MSL** | **Eye mask** | 18.46 | 18.83 | 17.51 – 19.42 | 0.48 |
|  | **Control mask** | 18.71 | 17.83 | 17.69 – 19.74 | 0.52 |
| **PAL,**  **Absolute**  **overnight change** | **Eye mask** | -1.32 | -1.00 | -2.13 - -0.52 | 0.40 |
|  | **Control mask** | -0.89 | -1.00 | -1.74 - -0.03 | 0.43 |
| **MSL,**  **Absolute**  **overnight change** | **Eye mask** | 1.20 | 1.50 | 0.62 – 1.77 | 0.29 |
|  | **Control mask** | 1.45 | 1.34 | 0.98 – 1.91 | 0.23 |
| **EXPERIMENT 2** | | | | | |
| **PAL** | **Eye mask** | 69.89 | 71.50 | 66.10 – 73.68 | 1.846 |
|  | **Control mask** | 67.71 | 71 | 63.76 – 71.67 | 1.927 |
| **PVT** | **Eye mask** | 319.64 | 319.55 | 303.88 – 335.40 | 7.705 |
|  | **Control mask** | 326.97 | 325.97 | 313.23 – 340.72 | 6.722 |

*Note:* PAL=Paired associate learning task. PVT= Psychomotor vigilance test. MSL=Motor-skill learning task. CI=confidence interval, SEM=standard error of the mean.

Table S2.

*LME models outputs, Experiment 1.*

| **Term** | **Estimate(SE)** | **95% CI** | **p** | **AIC**  **of a full model** | **AIC**  **of a reduced model** |
| --- | --- | --- | --- | --- | --- |
| **PAL** | | | | | |
| Intercept | 65.06(0.675) | 63.74 – 66.38 | <.001*** | 1026.0 | 1029.2 |
| Mask_type(0) | -1.19(0.514) | -2.20 – -0.19 | .020* |  |  |
| **PVT** | | | | | |
| Intercept | 316.37(3.723) | -0.11 – 0.28 | <.001*** | 3229.9 | 3235.2 |
| Mask_type(1) | -6.103(2.240) | -0.28 – -0.05 | .006** |  |  |
| **Absolute Overnight change - PAL** | | | | | |
| Intercept | -1.32(0.414) | -2.14 – -0.51 | <.001** | 910.77 | 909.61 |
| Mask_type(0) | 0.43(0.470) | -0.49 – 1.36 | 0.358 |  |  |
| **Absolute Overnight change - MSL** | | | | | |
| Intercept | 1.20(0.260) | 0.69 – 1.71 | <.001*** | 801.74 | 799.51 |
| Mask_type(0) | 0.25(0.348) | -0.43 – 0.93 | 0.478 |  |  |

*Note:* PAL=Paired associate learning task. PVT= Psychomotor vigilance test. MSL=Motor-skill learning task. Standard errors are given in parentheses. Predictor was coded as follows: ‘Mask_type’ – eye mask=1, control=0. AIC- Akaike information criterion. CI–confidence interval. ***p<.001; ** p<.01.

Table S3.

*Sleep parameters (mean±SEM) and pairwise comparisons.*

| **Sleep Measures** | **Eye Mask** | **Control Mask** |  | |  | |  |  |
| --- | --- | --- | --- | --- | --- | --- | --- | --- |
|  |  |  | **Statistical test** | **Test statistic** | | **p-value** | | **Effect Sizes** |
| TST (minutes) | 451.52±14.03 | 459.76±18.05 | t-test | -0.605 | | 0.550 | | 0.11 |
| Onset (minutes) | 15.83±2.11 | 14.72±2.60 | Wilcoxon | -0.470 | | 0.638 | | 0.08 |
| N2 (minutes) | 185.34±8.24 | 179.07±11.46 | t-test | 0.612 | | 0.545 | | 0.11 |
| N3 (minutes) | 99.10±6.18 | 107.96±6.48 | t-test | -1.610 | | 0.119 | | 0.29 |
| REM (minutes) | 123.00±10.20 | 133.96±7.92 | t-test | -1.143 | | 0.263 | | 0.21 |
| Wake (minutes) | 28.24±4.28 | 24.03±2.74 | Wilcoxon | -0.249 | | 0.804 | | 0.04 |
| No. awakenings | 2.14±0.39 | 2.03±0.32 | t-test | 0.351 | | 0.729 | | 0.06 |

*Note:* paired-samples t-test (t-test); Wilcoxon signed-rank test (Wilcoxon). TST=total sleep time; N2=non-rapid eye movement sleep stage2; N3= non-rapid eye movement sleep stage3; REM= rapid eye-movement sleep.

Table S4.

*LME models outputs, Experiment 2 and Experiment 1&2 combined.*

| **Term** | **Estimate(SE)** | **95% CI** | **p** | **AIC**  **of a full model** | **AIC**  **of a reduced model** |
| --- | --- | --- | --- | --- | --- |
| **EXPERIMENT 2** | | | | | |
| **PAL** | | | | | |
| Intercept | 67.714(1.853) | 64.08 – 71.35 | <.001*** | 389.09 | 391.00 |
| Mask_type(1) | 2.179(1.063) | 0.09 – 4.26 | .049* |  |  |
| **PAL – without N=4 outliers** | | | | | |
| Intercept | 69.769(1.199) | 67.42 – 72.12 | <.001*** | 330.56 | 333.31 |
| Mask_type(1) | 2.346(1.028) | 0.33 – 4.36 | .030* |  |  |
| **PVT** | | | | | |
| Intercept | 326.976(7.108) | 313.04 – 340.91 | <.001*** | 596.00 | 595.83 |
| Mask_type(1) | -7.34(5.343) | -17.81 – 3.14 | .18 |  |  |
| **EXPERIMENT 1 & 2** | | | | | |
| **PAL** | | | | | |
| Intercept | 65.13(1.220) | 62.74 – 67.53 | <.001*** | 1426.8 | 1433.8 |
| Mask_type(1) | 1.44(0.470) | 0.52 – 2.36 | .002** |  |  |

*Note:* PAL=Paired associate learning task. PVT= Psychomotor vigilance test. Standard errors are given in parentheses. Predictor was coded as follows: ‘Mask_type’ – eye mask=1, control=0. AIC- Akaike information criterion. CI–confidence interval. ***p<.001; **p<.01; *p<.05.
